# Engineered TasR genome editors enhance target-motif-independent editing in plants

**DOI:** 10.64898/2026.11.23.768706

**Authors:** Cynthia I. Terrace, Luke M. Oltrogge, Viktoriya Georgieva, Nicolas M. Ayala, Hanlun Jiang, Jennifer Listgarten, David F. Savage

## Abstract

TIGR-Tas systems are a new class of compact RNA-guided DNA endonucleases that are structurally and mechanistically distinct from CRISPR-Cas systems. Unlike CRISPR-Cas nucleases and other miniature RNA-guided endonucleases, which depend on restrictive target motifs such as PAMs or TAMs, TIGR-Tas effectors function without stringent motif requirements. Although an expanded target space is highly desirable, wild-type TasR enzymes exhibit very low editing activity, which limits their practical use. Here, we engineered ParTasR to enhance its activity for genome editing using functional yeast-based screens on diverse iterative libraries. We identified high-activity variants and evaluated them *in planta,* where precise editing is critical for quantitative trait engineering. The enhanced variants retained motif-independent targeting, preserving the broad targetable genomic space of the wild-type enzyme. Two engineered variants, TasRe4 and TasRe9, achieved editing efficiencies of up to approximately 23% at select loci *in planta*, a roughly 50-fold improvement over wild-type TasR. These results establish engineered TasR variants as promising, motif-independent genome-editing tools for plants.

**Significance:** Genome-editing tools that can precisely modify a wider range of DNA sequences could open new possibilities for improving crops. However, many existing tools, including CRISPR-Cas systems, can only modify DNA at sites next to a specific sequence, limiting where changes can be made. TIGR-Tas systems are compact genome-editing tools that can target DNA without requiring a specific target motif. Notably, the TasR ribonucleoprotein (RNP) is ∼25% the size of the SpCas9 RNP, potentially expanding delivery options for plant genome editing. However, wild-type TasR enzymes exhibit relatively low editing activity. Through iterative protein engineering, we developed two improved variants, TasRe4 and TasRe9, that edit plant DNA up to approximately 50 times more efficiently than the wild-type enzyme while retaining the broad target-site flexibility. These improved tools could enable modification of genomic regions difficult or impossible to reach with existing approaches, expanding opportunities for crop improvement and agricultural biotechnology.

## Introduction

**T**andem **I**nterspaced **G**uide **R**NA (TIGR) - **T**IGR-**as**sociated (Tas) systems are a new class of miniature RNA-guided DNA endonucleases that are mechanistically and structurally distinct from CRISPR–Cas systems ^1^. TIGR-Tas effectors universally contain an RNA-interacting nucleolar protein (Nop) domain but exhibit diverse architectures, including members that possess fused nuclease domains, such as RuvC and HNH in TasR and TasH effectors, respectively. These systems utilize guide RNAs (tigRNAs) characterized by a dual spacer array architecture, which enables coordinated targeting and cleavage of both strands of a target DNA sequence. TIGR-Tas effectors are highly compact, with a protein size of ∼300-400 amino acids (aa) and a mature tigRNA length of just 36 nts.

Other RNA-guided DNA endonucleases, including the widely used CRISPR-Cas systems, their evolutionary predecessors TnpBs and IscBs, and Fanzors, require highly restrictive target motifs such as Protospacer Adjacent Motifs (PAMs) and Transposon Adjacent Motifs (TAMs), respectively, to license DNA cleavage^2–6^. These motifs are typically 2-6 bps in length and consequently limit the targetable sequence space of a genome^7,8^. Engineering efforts have relaxed target motif sequence requirements across a variety of nucleases to expand targetable sequence space ^9–12^. However, this often comes at the cost of reduced cleavage activity, reflecting the role of these motifs in target search and cleavage kinetics^13^. TIGR-Tas endonucleases can operate without strict target-motif requirements, thereby expanding the range of nucleic acid sequences that may be targeted relative to other RNA-guided endonuclease platforms.

Given their apparent innate lack of target motif requirements, TIGR-Tas systems could be uniquely useful genome editing effectors. When constrained by target motif requirements, researchers must often use multiple endonucleases with distinct editing efficiencies, size constraints, and programming rules to access diverse target sites. This limitation is particularly consequential in plant biology where, for example, the editing of promoter cis-regulatory elements is increasingly being used for quantitative trait engineering^14–17^. Recent evidence indicates that single nucleotide resolution is critical for tuning the expression modulated by cis-regulatory elements, which can in turn have dramatic effects on plant architecture and yield^16,18^. However, many such prospective edit sites remain inaccessible to currently available genome-editing tools. TIGR-Tas endonucleases could help overcome these constraints by expanding the range of genomic sites accessible for precise editing.

### Engineering enhanced TIGR-Tas variants

Despite their target-motif-independent targeting capability, wild-type (WT) TasR enzymes exhibit extremely low editing activity, restricting their utility. Here, we used high-throughput functional screening on iterative libraries to engineer enhanced genome-editing activity in *Parcubacteria* TasR (ParTasR). Starting from an error-prone PCR library we used computational methods to nominate activating mutations expected to favorably combine and created a combinatorial library of stacked mutants, many of which greatly exceeded the activity of WT TasR. After testing a selection of high-activity variants in *N. benthamiana* we identified two engineered variants, TasRe4 and TasRe9, with substantially enhanced editing *in planta*, achieving editing efficiencies of up to approximately 23% at select genomic loci and representing roughly 50-fold improvements over wild-type TasR.

## Results

### In vivo activity-based selection in Saccharomyces cerevisiae

To assess on-target cleavage activity, we adapted an *in vivo* activity-based selection assay which leverages engineered yeast (*Saccharomyces cerevisiae*) strains with a genomic *ade2−* reporter cassette (Fig. 1)^19,20^. Under non-selective conditions supplemented with adenine, all transformed cells are maintained regardless of the cleavage activity of their respective TasR variant. In contrast, under selective conditions lacking adenine, only cells expressing TasR variants capable of on-target DNA cleavage undergo single-strand annealing (SSA)-mediated repair of the *ADE2* reporter, restoring adenine prototrophy and enabling cell growth. Thus, differential growth under selective and non-selective conditions provides a functional readout of target-site cleavage activity.

**Figure 1.**
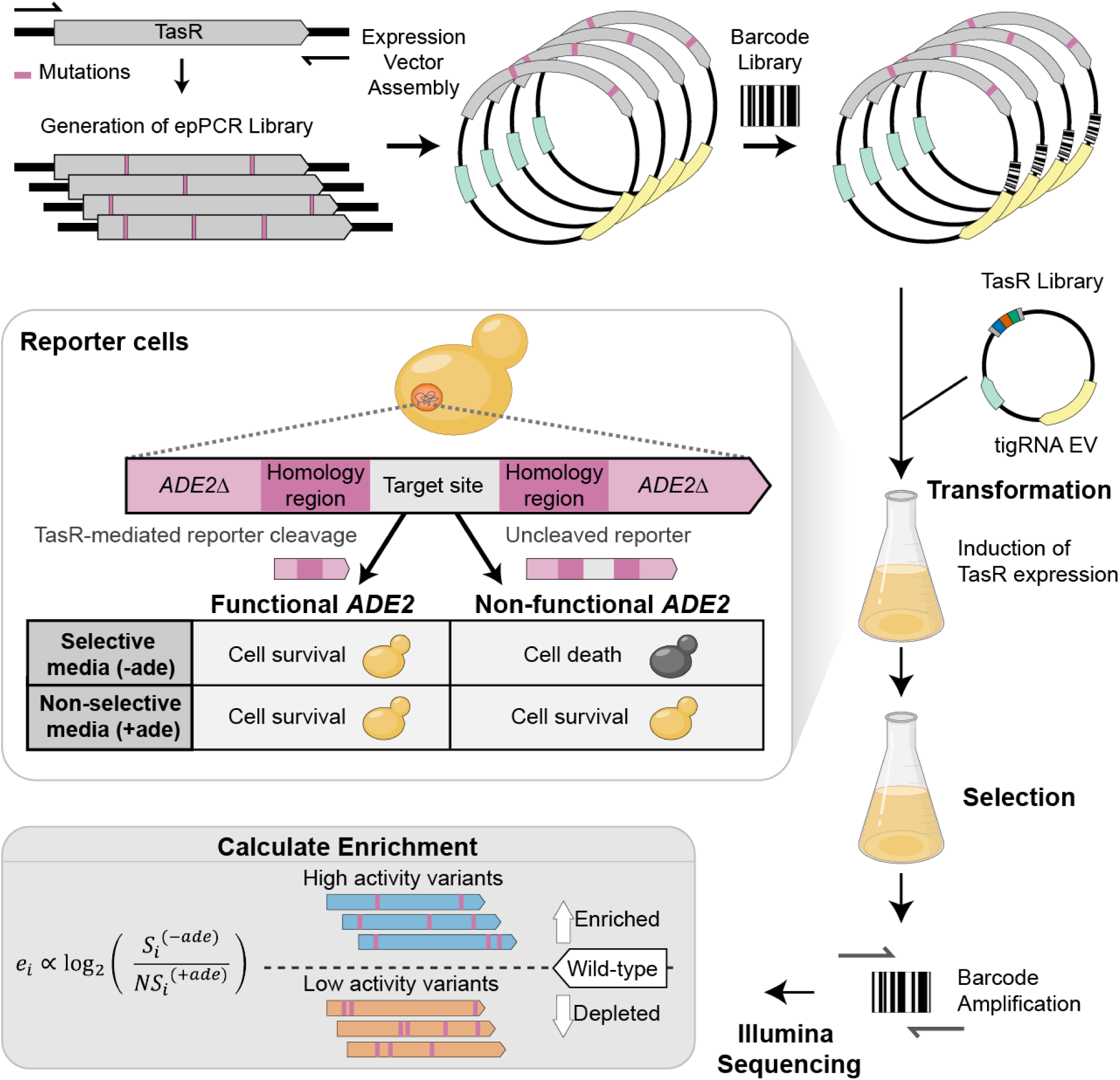
: Generation of a mutagenized TasR library and *in vivo* screening in yeast. Schematic of TasR error-prone PCR library construction, barcoding, and selection in yeast. On-target cleavage of the ADE2 reporter cassette initiates DNA repair, restoring *ADE2* function and enabling yeast growth in adenine-deficient media. Variant barcodes are sequenced, and enrichment calculated as the log ratio of barcode abundance under selective (-ade) and non-selective (+ade) conditions.

Among the previously characterized TasR orthologs, ParTasR demonstrated the highest editing activity in mammalian cells^1^ and was therefore selected for the engineering campaign. The highly-compact 337 amino acid ParTasR protein sequence, codon-optimized for expression in *S. cerevisiae* and human cells, was designed for assembly into a yeast expression vector with nuclear localization sequence (NLS) fused to the carboxyl terminus (C-terminus).

Using error-prone PCR (epPCR), we generated a pooled plasmid library of ∼ 1.5 x10^5^ TasR protein variants with an average mutation rate of 3.2 mutations per variant. The library was barcoded to enable downstream variant-level analysis, and variant-barcode associations were established through long-read sequencing (See Methods and *SI Appendix*, Table S1).

### epPCR Library reveals activating hotspots

The epPCR library and the wild-type tigRNA counterpart were transformed into the yeast reporter strain. After inducing expression for 8 hours, the yeast libraries were grown out under selective (-ade) and nonselective (+ade) conditions with two biological replicates. Variant barcode abundance was quantified with short-read next-generation sequencing (NGS) across all conditions. Barcode enrichment was calculated as the log ratio of selective over non-selective barcode abundance. These scores were then aggregated to variant-level enrichment by taking the median across corresponding barcodes (Fig. 2A). To place all variants on a common functional scale, we defined two internal reference classes: nonfunctional (dead) controls comprising variants that introduce a stop codon, and wild-type controls comprising synonymous mutation variants that preserve the wild-type protein sequence. Only control variants with ≥10 reads in both replicates were used. Enrichment values were then normalized so that the dead median corresponds to 0 and the wild-type median to 1.

**Figure 2:**
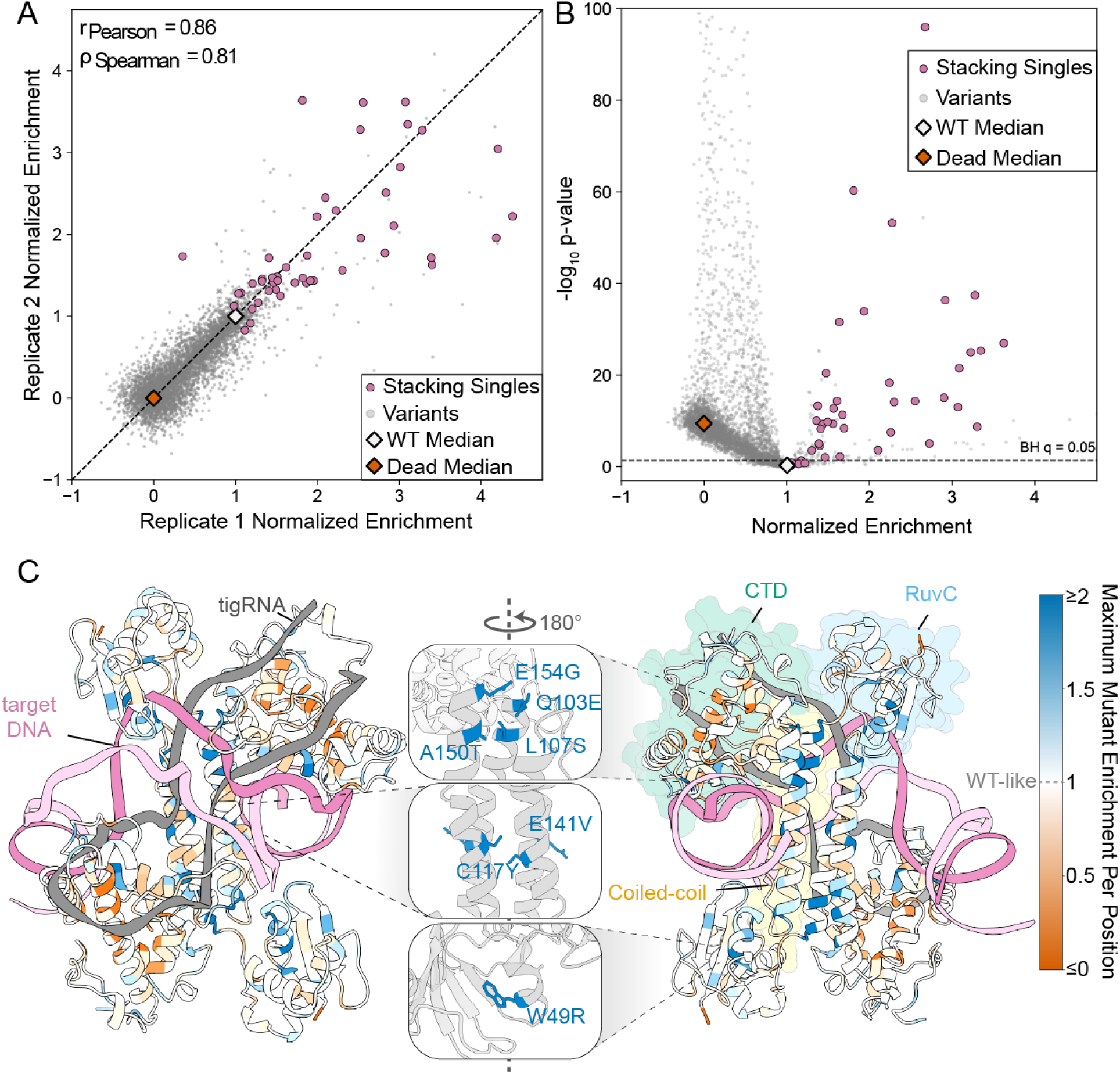
Error-prone PCR library reveals activating hotspots. (A) The normalized enrichment scores of epPCR library variants across two experimental replicates at target site 1. Variants with ≥10 barcodes across the two replicates are mapped in grey. Single mutation variants (Stacking singles) across the selected 33 residue positions for second-generation stacking libraries shown in pink. The WT Median control is the median enrichment value of all protein variants encoding the wild-type protein sequences via synonymous mutations displayed as an outlined white diamond. The Dead Median control represents the median enrichment value of variants containing premature stop codons displayed as a vermillion colored diamond. (B) Volcano plot showing mean enrichment of variants and corresponding statistical significance following selection (n = 5,725 variants with ≥10 barcodes in each of two replicates; stop-codon and synonymous variants excluded except as displayed averages). Statistical significance was calculated using individual barcode measurements with a two-sided Mann–Whitney U test with the Benjamini-Hochberg multiple hypothesis correction. The WT and Dead medians are indicated by outlined white and vermillion diamonds, respectively. Single-mutant variants at the 33 selected residue positions used for second-generation combinatorial library construction are highlighted in pink. (C) AlphaFold ParTasR model colored by positional maximum normalized enrichment scores aligned to the nucleic acid of the TaTasR ribonucleoprotein (RNP) Cryo-EM (9MTY). The tigRNA is shown in grey, and the target DNA strands are shown in pink. ParTasR domain annotations highlight a single protein monomer within the dimeric structure. Close up views of enriched mutations displayed in the center of the two structural views that differ by 180°.

The epPCR library captured 99.9% of all possible single nucleotide mutations. However, given that many specific amino acid mutations require multiple nucleotide mutations to reach, only a subset are present in the data. In total, 30.8% (1,977 of 6,422) of possible single-amino-acid substitutions were represented in the library (*SI Appendix*, Fig. S1A). The library also captured substantial combinatorial diversity, including 2.1x10^4^ double-mutant variants, 1.4x10^4^ triple-mutant variants, and 1.2x 10^4^ higher-order variants (>3 mutations).

Of the observed single-amino-acid substitutions, 3.3% had enrichments significantly above wild-type (Benjamini–Hochberg-adjusted *p* < 0.05; Fig. 2B). These mutational hotspots were shown to cluster in the tertiary structure when the maximum enrichments were plotted onto the AlphaFold model (Fig. 2C and *SI Appendix*, Fig. S1B). Two prominent clusters of activating mutations localized to the RuvC nuclease domain and coiled-coiled dimerization domain.

Among these enriched positions, residue 49 within the RuvC nuclease domain exhibited a preference for non-wild-type substitutions, with the wild-type tryptophan (W) consistently disfavored relative to all other observed substitutions at this position. To further assess the activating substitutions identified in the pooled library screen, we selected individual variants for experimental validation. These single-mutant variants exhibited increased activity relative to wild-type TasR (*SI Appendix*, Fig. S2F).

### Second-generation combinatorial library yields enhanced protein variants

To explore increases in TasR editing activity through combinations of single-mutant variants, we selected 35 positions across the TasR protein sequence (Fig. 2A and Methods). The initial epPCR library screen enabled quantitative assessment of variant activity, revealing activating single-amino-acid substitutions distributed throughout the TasR protein sequence Using the selection data of the initial library, we trained a neural network to predict the enrichment of pairwise combinations of activating single substitutions. From these activating double mutations predicted by our model, a set of mutually compatible single substitutions were selected to generate a second-generation combinatorial library. This included 33 positions targeted for missense mutations and 2 positions targeted for synonymous mutations as internal wild-type controls. Additionally, because W49 had been identified as a key position, we incorporated an NNN codon at W49 to explore alternative substitutions while generating premature stop codons at a low frequency that could serve as an internal dead control.

We reasoned that many activating mutations may be slightly destabilizing on average and the accumulated effect can be a loss of folding free energy, ultimately compromising proper folding^21^. To mitigate this possibility, we included 3 mutations (L15W, A46S, E152Q) that exhibited the highest median enrichment among the double-mutant variants but were not included in the original set of nominated mutations. Notably, these substitutions were observed to rescue activity when combined with otherwise deleterious mutations. We postulated that this compensatory behavior may reflect a protein-stabilizing effect, potentially promoting proper folding in the context of multiple activating mutations. Together, these designs encompassed 65 possible amino acid mutations for the second-generation combinatorial library.

Using a nicking mutagenesis method, we constructed a combinatorial library of ∼1.34 x10^4^ variants with an average of 4.15 mutations per variant (Fig. 3A)^22,23^. We barcoded the library such that there was a median of 24 unique barcodes per variant.

**Figure 3:**
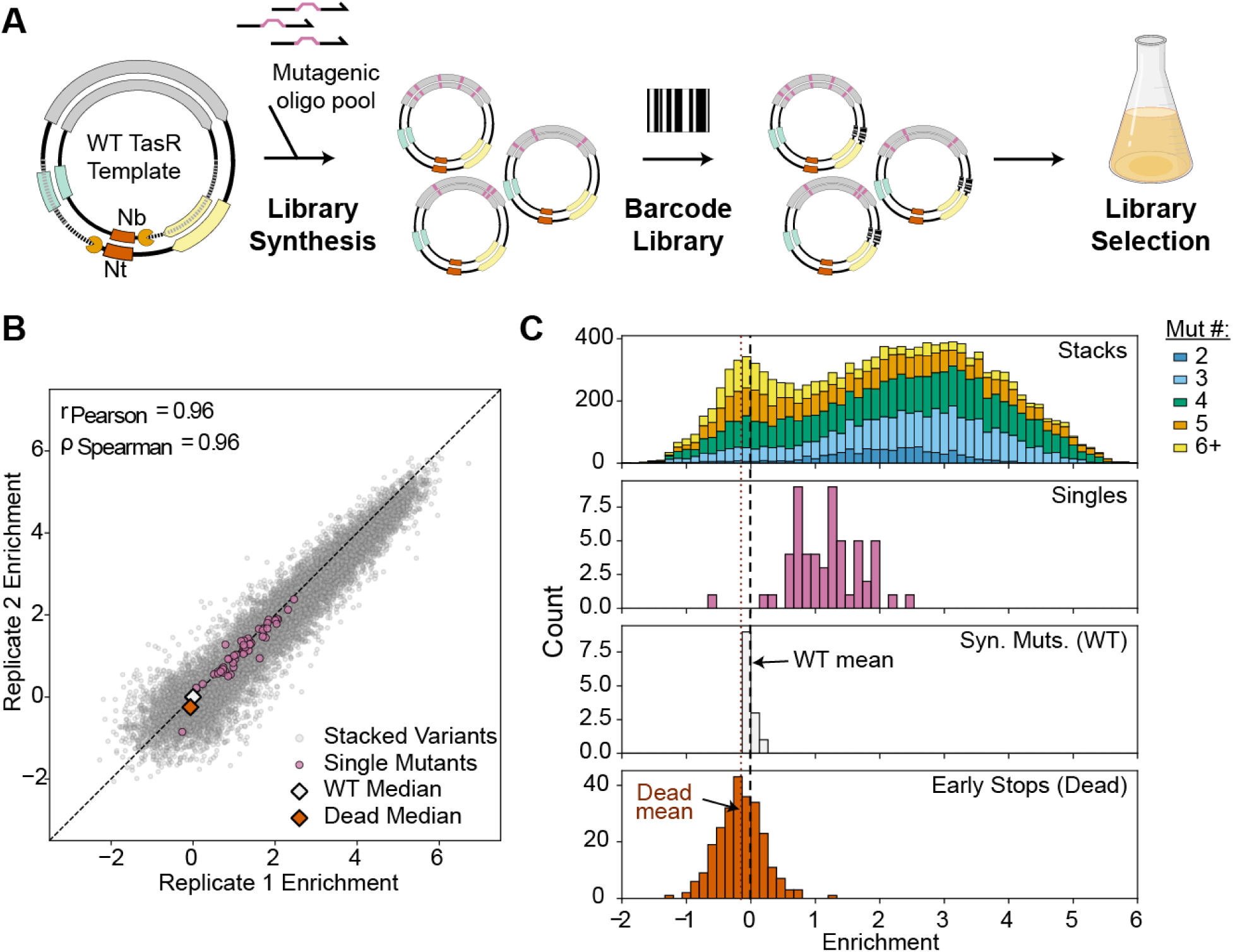
Second-generation combinatorial library yields enhanced protein variants. (A) Schematic representation of the second-generation stacked mutation library construction built via nicking mutagenesis and subsequent screening at target site 1. (B) Enrichments of the stacked library across two experimental replicates showing variants with ≥5 barcodes. Due to the high average activity of this library the WT (WT synonymous) and dead (premature stops) variants have similar raw enrichments. The enrichments are therefore normalized just to WT such that its median enrichment is 0. Single mutation variants across the selected 33 residue positions for second-generation stacking libraries (shown in pink). (C) Histograms showing the distribution of variant activity by mutation class including: stacks of two or more mutations, the observed single mutants, WT synonymous, and premature stop codons (i.e. “dead”). Only variants represented by ≥5 barcodes in each replicate were included.

The combinatorial library was screened in the same yeast reporter assay as described above, and exhibited robust activity, with many variants displaying substantially higher enrichments than either wild-type TasR or any of the parent single-mutant variants (Fig. 3B). The distribution of the library composition across the activity range demonstrates that many of the nominated single mutations combined synergistically to generate a diverse population of functional combinatorial variants. Even among the most active variants, higher-order mutation combinations were prevalent, with a substantial fraction of the population harboring 4-5 mutations (Fig. 3C).

As the library screens were conducted against a single target sequence, we wanted to ensure that variants had no protein-dependent sequence specificity and would generalize to other target sites. We assembled a 50-member variant library for screening at an alternative target site (*SI Appendix*, Fig S2A, S2B, S2C). This hit-derived mini-library comprised 48 variants identified from the second-generation combinatorial screen, including 38 of the highest activity variants and 10 additional variants selected to span the observed activity range and serve as internal controls for benchmarking relative activity. Wild-type TasR and catalytically inactive (D7A) dTasR, were included as additional positive and negative controls, respectively.

The rank order of variant activity was generally preserved at the secondary target site. Notably, while the top 38 variants exhibited largely indistinguishable activity at the initial target site, screening at the secondary target site resolved sequence-divergent, high-performing variants (*SI Appendix*, S2C, S2E). Select variants exhibiting high activity at both target sites in yeast and diverse mutation profiles were selected for individual characterization yielding substantial increases in activity relative to wild-type TasR, with activity levels also markedly exceeding those observed for the single-mutation variants (*SI Appendix*, Fig. S2).

### Enhanced combinatorial TasR variants screened in *Nicotiana benthamiana* via agroinfiltration

To further evaluate the activity of the combinatorial enhanced TasR variants, six enhanced variants were subsequently assessed in the model dicot *Nicotiana benthamiana* at three genomic loci. The selected variants were delivered by agroinfiltration into *N. benthamiana* leaves, and indel frequencies of infiltrated tissue were quantified by NGS using CRISPResso (version 2.3.3) (Fig 4A). Wild-type TasR displayed low, locus-dependent editing activity across all tested loci. The combinatorial variants outperformed the wild-type TasR across all tested loci, with two variants, TasRe4 (G129R; V148I; E216K; A233S; T312A) and TasRe9 (L15W; E141V; A150T; T312A) exhibiting high activity at every locus (Fig 4B). Interestingly, both enhanced TasR variants generated a diverse range of repair outcomes, with deletions representing the predominant outcome and spanning a broad size range of 2 to 40+ bp (*SI Appendix*, Fig. S3 and Fig. S4).

**Figure 4:**
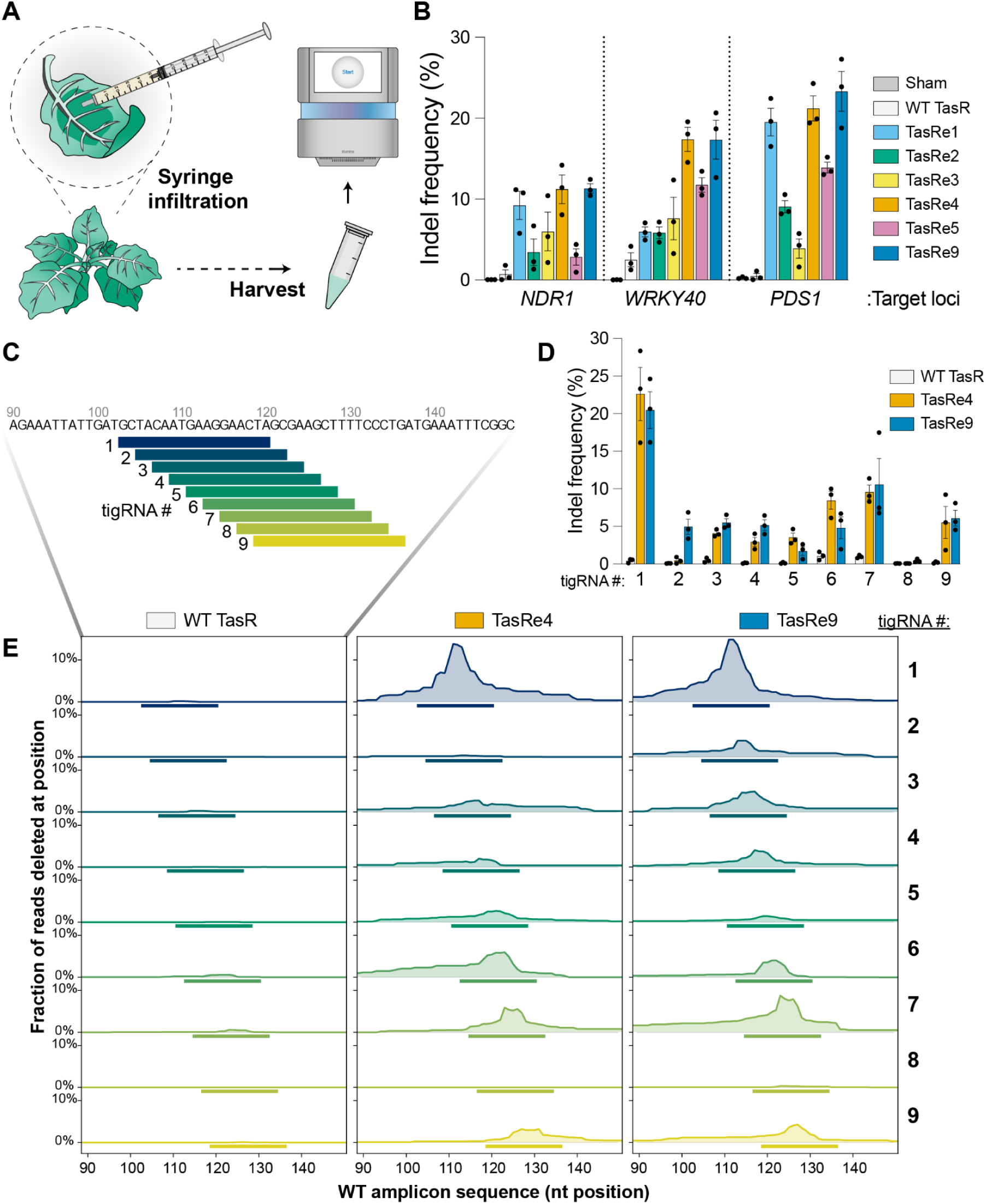
**Engineered TasR variants exhibit robust PAM-less editing *in planta.*** (A) Schematic representation of *N. benthamiana* transformation via syringe infiltration of agrobacterium carrying TasR RNP expression vector. (B) Indel frequencies of six of the highest-activity stacked library variants TasR variants across three genomic loci in *N. benthamiana*. Data from independent biological replicates (n =3) are plotted as the mean ± s.e.m. (C) Schematic illustrating nine tiled tigRNAs within the shown 60 bp window of the Nb*PDS1* locus. (D) Overall indel frequencies of WT, TasRe4, and TasRe9 across the nine tiled guides targeting the Nb*PDS1* locus, with each successive guide shifted 2 bp downstream. Data represent the mean ± s.e.m. of three independent biological replicates (n = 3). Background correction was performed by sham subtraction: for each indel signature, the mean frequency in the sham (buffer-only) control was subtracted from the corresponding sample frequency, with negative values set to zero. (E) Per-position deletion profiles across the Nb*PDS1* target site for nine tiled guides evaluated with wild-type (WT), TasRe4, and TasRe9. Deletion frequencies are shown across a 60 bp window of the WT amplicon for WT TasR (left), TasRe4 (middle), and TasRe9 (right). Each trace represents a single guide, with the nine guides (1-9) arranged from top to bottom in tiling order. The bar beneath each trace denotes the corresponding 18 bp target sequence. At each position, the trace indicates the fraction of aligned reads containing a deletion spanning that position. Values represent the mean of three biological replicates.

### TasRe4 and TasRe9 are enhanced miniature editors without target motif constraints

Unlike other programmable RNA-guided DNA endonucleases, whose targetable sequence space is constrained by target motif requirements, TasR endonucleases are not subjected to such constraints. To evaluate this expanded targeting potential, the two highest-performing variants, TasRe4 and TasRe9, were targeted to the Nb*PDS1* locus using a series of tiling guides shifted downstream in 2 bp increments, yielding nine target sites spanning the 18 bp target sequence (Fig. 4C, 4D, 4E).

Wild-type TasR exhibits consistently low editing efficiencies (≤1%) across target sites, whereas the engineered variants achieve substantially higher editing efficiencies of ∼ 5-22% at eight of the nine target sites (Fig. 4D). TasRe4 and TasRe9 exhibit comparable editing efficiencies across most sites; however, at site 2, TasRe9 achieves an approximately 11-fold improvement over TasRe4 and a 57-fold improvement over wild-type TasR. Both engineered variants and wild-type TasR exhibit low editing efficiencies at site 8, suggesting that this target site is less permissive to editing. Indel profiles of the engineered variants reveal a shifting window of repair that tracks with the repositioning of the tigRNA target window, whereas wild-type TasR exhibits nearly undetectable indel frequencies across all nine sites (Fig. 4E). Although editing efficiencies varied across the tiled guides, indicating some sequence context dependence, the engineered variants retain target-motif-independent editing activity across multiple target sites, broadening the targetable genomic space and enabling editing at previously inaccessible positions.

## Discussion

In this study, we leveraged iterative functional screening and complementary library design strategies to engineer two highly active TasR variants, TasRe4 and TasRe9. These variants achieved editing efficiencies of up to ∼23% in *N. benthamiana*, representing a roughly 50-fold improvement over wild-type TasR. By combining selected substitutions in a second-generation library, we demonstrated that activating, single-amino-acid substitutions identified through an epPCR screen could be combined to generate higher-order TasR variants with enhanced activity. Notably, the engineered variants retain the target-motif-independent targeting capability of wild-type TasR, preserving its broad targetable genomic space while substantially improving editing efficiency across tested sites. The high-activity variants identified in these screens harbor diverse mutation profiles with limited overlap, suggesting there is additional room for improving TasR activity.

The abundance of activating mutations identified in TasR mirrors observations for the transposon-associated endonuclease TnpB, suggesting that, like TnpB, native TasR is not optimized for high cleavage activity and may retain substantial capacity for improvement^19^. Our results identify a diverse set of activity-enhancing mutations, highlighting multiple routes to increased editing activity. However, the relatively limited structural information available for TasR, coupled with the dynamic and numerous conformational states that RNA-guided endonucleases adopt during target search and recognition, makes it difficult to infer the mechanistic contributions of these mutations or assign specific functional roles to them across individual structural states^24,25^. Future biochemical studies may elucidate how these activating mutations influence RNP stability and catalytic function.

Realizing the full potential of TIGR-Tas systems as genome-editing tools will require further characterization of their targeting specificity. Unlike PAM-dependent CRISPR-Cas systems, for which numerous tools have been developed to characterize and predict off-target editing, TIGR-Tas systems lack comparable predictive frameworks. Because the target-search mechanisms of these two enzyme classes likely differ, existing CRISPR-Cas prediction tools may not accurately capture TIGR-Tas off-target activity^26,27^. Developing reliable methods to characterize and predict the on-and off-target activity profiles of TasR will therefore be an important next step. Further investigation into the sequence features and design principles governing tigRNA function may help inform the development of these predictive approaches. In particular, understanding the determinants of target recognition will be important for evaluating the potential for off-target activity associated with TasR’s motif-independent targeting capability.

PAM requirements constrain target recognition in CRISPR-Cas systems and thereby limit off-target editing, whereas PAM-relaxed variants have been associated with increased off-target activity^24^. Because TasR variants lack a requirement for a PAM or TAM to license target cleavage, they may similarly exhibit an increased propensity for off-target editing. In plants, however, this potential risk can be mitigated through phenotypic screening of regenerated individuals, complemented by whole-genome sequencing to identify and cull lines with unintended genomic alterations.

Future efforts integrating machine learning-guided protein engineering with improved characterization of target specificity and tigRNA design principles may further advance editing performance and expand the utility of these compact genome editors. Collectively, these engineered TasR variants represent promising tools for genome editing applications in agriculture and biotechnology.

## Methods

### Error prone PCR library construction

The ParTasR protein sequence was codon optimized for expression in *S. cerevisiae* and mammalian cells and purchased from Twist Bioscience as a gene fragment with flanking restriction enzyme recognition sites for assembly with an intermediate vector by Golden Gate cloning with BsaI-HF.

The error prone PCR library (epPCR) was constructed using the GeneMorph II Random Mutagenesis Kit (Agilent Technologies) with 100 ng template input. Cycling conditions included an initial denaturation at 95°C for 2 minutes, 30 cycles of 95°C for 30 seconds, 62°C for 30 seconds, and 72°C for 1 minute and 45 seconds, and a final extension at 72°C for 10 minutes.

The library was cloned into a destination vector by Golden Gate cloning with Esp3I.

The epPCR library was digested with NotI-HF and XhoI and barcoded by assembly (NEBuilder HiFi DNA Assembly Master Mix) with ssDNA oligonucleotides with internal 30xN barcodes. The barcoded plasmid assembly was transformed into Top10 Electrocompetent *E. coli* and bottlenecked such that a larger culture for plasmid purification was inoculated with ∼7.1x10^5^ transformed cells, estimated from colony-forming units (CFUs) counted from titer plates.

### *In silico* selection of mutations for stacked library

To select the candidate substitutions to be stacked in the oligo pool nicking mutagenesis library, we considered the predicted enrichment of pairwise combinations of the most activating single substitution mutations. For the 207 single substitutions which were enriched over wild-type ParTasR there are 21,321 possible pairwise combinations, of which only 118 were directly observed in the initial epPCR library selection. To predict the log-enrichment of the unobserved double mutations, we trained a neural network whose input features include ESMC representations of the double mutant and its constituent single mutants, as well as the measured log-enrichment of both single mutants ^28^. The neural network first takes the three ESMC representations and projects each residue embedding down to 8 dimensions. The three representations are then flattened into a single vector, and passed through a two-layer multilayer perceptron (MLP) with Gaussian error linear unit (GELU) activations and a hidden dimension of 256. The MLP output, together with the log-enrichments of the two single mutations, are then passed through two more MLP layers which produce the log-enrichment of the double mutant. We used this neural network to predict the log-enrichment of all possible pairwise combinations of the 207 activating substitutions, and selected a subset of 50 substitutions whose pairwise combinations within the set had the highest average predicted log-enrichment.

The selection was performed by a greedy algorithm, where we started from an initial set of candidate single substitutions, and swapped substitutions into the candidate set which gave the highest improvement to the resulting predicted average log-enrichment of pairwise combinations. Specifically, using the predictions of the pairwise combinations of the 207 activating substitutions, we constructed a 207 by 207 symmetric matrix where each row or column index corresponded to an activating substitution. Each element e_ij_ is the predicted log-enrichment of the double mutation combining the single mutation i with the single mutation j. Elements whose indices were mutations on the same position were ignored, as those double mutations were not possible. We selected a set of 50 indices which approximately maximized the sum of their corresponding submatrix. To achieve this, the algorithm started with a set of the 50 most activating substitutions and performed a series of index swaps. Each swap took the index from within the set that contributed least to the submatrix sum, and swapped it with the index outside the set which would contribute the most to the submatrix if it were included. The swaps were continued until we could no longer improve the sum of the submatrix.

### Stacked library construction

Two epPCR library datasets were generated using slightly different tigRNA expression designs and selection conditions. The *in silico* mutation nomination campaign was conducted on the first dataset while the second dataset was used for all data presentation since it is experimentally identical to the process for the succeeding stacked library. The two data sets were closely concordant with ρ Spearman = 0.77 for variants with ≥10 barcodes. Upon integrating data from both rounds of library screening, three mutations from the initially nominated set (Y18H, I187S, and E308G) exhibited lower pairing scores in the secondary dataset and were therefore omitted from the subsequent mutation pool. Conversely, three alternative mutations (L15W, A46S, and E152Q), which exhibited the highest median effects among double-mutant variants in the secondary dataset, were incorporated. Notably, these substitutions rescued activity when paired with otherwise deleterious mutations. We hypothesized that this compensatory behavior may reflect a protein-stabilizing effect, potentially promoting proper folding in the context of multiple activating mutations.

Beyond the initially nominated set of mutations, an NNN codon was introduced at position W49 to explore previously uncharacterized amino acid substitutions at this particularly interesting position, where the wild-type tryptophan appeared to suppress activity. The NNN codon also provided an opportunity to introduce premature stop codons, which served as an internal catalytically inactive control within the library.

The combinatorial library was constructed using nicking mutagenesis, which enabled the incorporation of multiple precisely specified mutations through the addition of mutation-containing single-stranded DNA (ssDNA) oligonucleotides ^22,23^. DNA oligos covering 35 residues with 65 possible mutations in the TasR protein sequence were phosphorylated and pooled in an equimolar ratio (*SI Appendix*,Table S2). Of the 35 mutated positions, two were included to induce synonymous mutations and act as internal wild-type controls. Due to the nature of the synthesis method, co-occurance of adjacent mutations is unlikely. To overcome this limitation and explore the outcomes of specific paired mutations, a set of dual mutation oligos were also added to the pool.

During the second strand synthesis, 50 pmol of the phosphorylated oligo was initially added to the reaction with 0.38 fmol of ssDNA template. Following five and ten cycles of amplification, an additional 43 pmol of the phosphorylated oligo was spiked into the reaction. The resulting library consisted of 13,374 variants and was barcoded such that there was a median of 24 unique barcodes per variant as measured via long-read sequencing.

### Hit-derived variant library construction

Selected variants were amplified using DNA from the combinatorial library yeast minipreps as the template, a shared forward primer, and a uniquely mapped barcode sequence as the reverse primer. Purified amplicons were then used as templates for a secondary PCR to add Golden Gate overhangs and Type IIS restriction enzyme recognition sites, facilitating cloning into a yeast expression plasmid. Wild-type and dead (D7A) protein sequences were amplified in a similar manner. The amplicons were subsequently purified, quantified using a NanoDrop, and pooled at equimolar concentrations. The resulting 50-member library was then barcoded such that each member was represented by a median of ∼17 unique barcodes.

### Yeast reporter strain generation

We used yeast reporter strains originally constructed and characterized by Thornton & Weissman et al. 2026. Two reporter strains with distinct target sites (Site 1 & Site 2) were used for subsequent library screening (*SI Appendix*, Table S3).

### Yeast-based activity assay for pooled library screening

We adapted an *in vivo* activity-based selection assay which leveraged engineered yeast reporter strains with a genomic *ade2−* reporter cassette^19,20^. A key modification here was the implementation of a liquid-based selection approach, which overcame limitations of the previous plating-based method. Specifically, this approach eliminates the need for pre-screen reversion-rate measurements and plating calibration, avoids the exclusion of low-activity enzymes imposed by reversion thresholds, and substantially reduces labor-intensive plate processing.

Prior to transformation, plasmid DNA was linearized with PaqCI restriction enzyme digestion. Linearized TasR plasmid libraries and tigRNA plasmids were transformed in a 1: 4 molar ratio. For the epPCR library, the second-generation combinatorial library, and the hit-derived variant library, 35µg, 14µg, and 1.2µg of linearized plasmid DNA, respectively, were transformed per experimental replicate. Yeast were transformed using the lithium acetate and single-stranded carrier DNA/PEG method^29^.

After transformation, expression vectors were assembled from the linear plasmid DNA by *in situ* gap repair homologous recombination in yeast, enabling growth in selective media. Transformed cells were resuspended in synthetic complete dropout medium (SCD) lacking leucine to select for transformation and recovered overnight at 30°C. The next morning, the cultures were induced at an initial optical density at 600 nm (OD_600_) of 1.0 with liquid medium lacking leucine with 2% galactose (w/v). Induced cultures were grown at 30°C, and samples were collected at multiple timepoints. Samples were pelleted, washed with Milli-Q water, and seeded into selective (SCD -adenine -leucine + 2% glucose (w/v)) and nonselective (SCD +adenine -leucine + 2% glucose (w/v)) liquid media at an initial OD_600_ of ∼ 0.5. Cultures were grown at 30°C for 48 h, after which samples were pelleted, and plasmid DNA was extracted using Zymoprep yeast plasmid miniprep II (Zymo Research).

### Yeast titer-based assessment of cleavage activity

To assess the activity of individual TasR variants, 0.5µg of linearized DNA encoding the TasR protein and tigRNA was transformed, assembled, and induced as described above. Induced cultures were grown at 30°C for 24 h, unless otherwise specified. At pre-induction and post-induction timepoints, approximately 1-2 OD_600_ units were removed from the transformed yeast culture, washed twice with Milli-Q water, and resuspended in 200µl Milli-Q water. Serial dilutions were prepared in triplicate and plated on selective (SCD -adenine -leucine + 2% glucose (w/v)) and nonselective (SCD +adenine -leucine + 2% glucose (w/v)) media. Plates were incubated at 30°C for 48 h. Colony reversion was used as a rough proxy for editor activity.

Colony reversion rates were calculated as the number of CFUs on selective medium divided by the number of CFUs on nonselective medium, multiplied by 100 to obtain the percentage.

Colony reversion percentages were plotted as bar graphs using GraphPad Prism (version 10.1.1).

### DNA sequencing

For the epPCR library and the second-generation combinatorial library, protein variant barcodes were amplified from plasmid DNA using KAPA HiFi HotStart ReadyMix (Roche) with 15 cycles for PCR1 and 10 cycles for PCR2. PCRs were purified with Ampure XP beads (Beckman Coulter) and submitted for 150-bp paired-end sequencing on Illumina NovaSeq sequencer at the UC Berkeley QB3 Genomics sequencing core (*QB3 Genomics, UC Berkeley, Berkeley, CA, RRID:SCR_022170)* and the Illumina NextSeq sequencer at the Innovative Genomics Institute (IGI) NGS sequencing core.

For the hit-derived variant library the protein sequence and barcoded region were co-amplified from plasmid DNA using KAPA HiFi HotStart ReadyMix (Roche) with 20 cycles. PCRs were cleaned with Zymo DNA Clean and Concentrator-5 (Zymo Research) and sent for native-ligation barcoding and nanopore sequencing (MinION Oxford Nanopore) at the UC Berkeley DNA Sequencing Facility.

### Variant-barcode mapping

After library construction, variants were associated with their barcodes using long-read sequencing (PacBio Revo 25M) at the UC Berkeley QB3 Genomics sequencing core. The reads were converted to FASTQ format with samtools (v1.19), mapped to the wild-type reference plasmid sequence with an N-masked 30-mer barcode using minimap2 (v2.28), and sorted to an indexed BAM file with samtools. Pysam was used to extract barcode and TasR ORF sequences. These were clustered by common barcodes, and the associated ORF variant consensus sequences were decided position-by-position as the majority base call. Only barcodes/variants with three or more reads were considered.

### Variant enrichment calculations

Barcode abundance in selective (S) and non-selective (NS) cultures was quantified from Illumina shortread sequencing using custom code. The 30-nt random barcode was extracted with cutadapt (v5.0) as a linked adapter built from the 20 nt of plasmid sequence immediately flanking the barcode on each side (max error rate 0.05, min overlap 10 nt). Reads lacking either flank were discarded and those having a barcode of exactly 30 nt were tallied for exact matches across each sample. Only barcodes having at least 6 counts in one or more NS samples were retained for enrichment analysis. For each replicate the enrichment for the ith barcode was computed as,

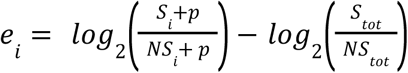

where *S_i_*and *NS_i_* are the reads counts in the paired selective and non-selective samples, *S_tot_* and *NS_tot_* are the total read counts, and *p* is the pseudocount constant (used as *p*=1) in our analysis.

Barcodes were associated to variants via exact matches to the longread mapping described above. Variant-wise enrichments were calculated as the median of all associated barcodes. In cases where normalization was applied, the variant enrichments were scaled so that premature stop codons (presumptively non-functional) had an average enrichment of zero while synonymous wild-type mutants had an average enrichment of one. Specifically, the normalized enrichment for the j^th^ variant was calculated as,

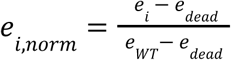

### Nicotiana benthamiana infiltrations

Plasmid DNA containing TasR protein and tigRNA expression cassettes were transformed into *Agrobacterium tumefaciens* strain GV3101 which was subsequently infiltrated into *N*. *benthamiana*.

Transformed *Agrobacterium tumefaciens* competent cells were grown on lysogeny broth (LB) agar medium supplemented with spectinomycin (50 µg ml^−1^), gentamicin (30 µg ml^−1^) and rifampicin (25 µg ml^−1^) at 30°C for 48 h. For each construct, a single, transformed colony was picked and inoculated into lysogeny broth (LB) medium supplemented with spectinomycin (50 µg ml^−1^), gentamicin (30 µg ml^−1^) and rifampicin (25 µg ml^−1^) and grown at 30°C overnight. The following morning, cultures were collected by centrifugation at 3,500 g for 10 min and resuspended in infiltration buffer (10 mM MgCl2, 10 mM MES (pH 5.6), and 150 µM acetosyringone, in Milli-Q water) to an OD_600_ of 1.0 and incubated at room temperature for 3 h. Using a 1 ml needle-less syringe, the prepared inoculum was syringe-infiltrated on the abaxial surface of three diagonally opposite leaves of 4-week-old *N. benthamiana* plants. Infiltrated plants were watered and recovered in the plant growth chamber (16-h light, 8-h dark photoperiod, 80 µmol m^−2^ light intensity, 50% humidity, at 23 °C) for 4 days.

To evaluate editing frequency of agrobacterium-infiltrated leaves, 4 days post-infiltration, 5 mm leaf disks were harvested from infiltrated areas. Harvested leaf tissue was resuspended in 100 µl of Phire Plant Direct PCR Dilution Buffer (Thermo Scientific, F160S), crushed with a 1 ml pipette tip and frozen to assist in lysis. Upon sample thawing, sample tubes were centrifuged for 5 min at 21,000 g, and the supernatant was transferred to a fresh Eppendorf tube. To amplify the region surrounding the target site, 2 µl of leaf lysate was used as template for the PCR reaction. PCR reactions were performed using the Phire Plant Direct PCR Master Mix (Thermo Fischer), and PCR products were cleaned up with Ampure XP beads (Beckman Coulter). Amplicons were then analyzed by a 4150 TapeStation (Agilent) and sequenced with 300-bp paired-end sequencing on Illumina NextSeq sequencer at the IGI. Editing frequency of the submitted samples were assessed by CRISPResso (version 2.3.3). To account for any systematic sequencing errors, background correction was performed by sham subtraction (Fig. 4D, 4E), in which the mean frequency of each indel signature in the sham (buffer-only) control was subtracted from the corresponding sample frequency. Negative values following subtraction were set to zero.

## Author contributions

C.I.T., L.M.O., and D.F.S. conceptualized the project. C.I.T. designed and conducted the experiments with assistance from L.M.O. and V.G.. C.I.T. and V.G. optimized the yeast assay.

H.J. and N.M.A. nominated variants for second generation library synthesis. C.I.T. and L.M.O. analyzed the data. C.I.T. and V.G. collected data on individual TasR variants in yeast and *N*. *benthamiana*. D.F.S. and J.L. supervised the study. C.I.T. and D.F.S wrote the manuscript with input from all authors. All authors reviewed the manuscript and approved the final version.

## Classification

Biological Sciences; Plant Biology

## Acknowledgements

We thank B.W. Thornton, R. F. Weissman, and J. E. Rodriguez for general guidance, establishing protocols and providing components of expression vectors that we used in the yeast cleavage assay and *N*. *benthamiana* infiltrations. We thank J. L. Rivera for cultivating the *N*. *benthamiana* plants used in this study. We thank N.G. Karavolias for insightful discussions regarding the potential agricultural applications of this work. We thank N. Krishnappa for assistance with running NGS samples at the IGI NGS sequencing core. This material is based upon work supported by the National Science Foundation graduate research fellowship program under grant no. DGE 2146752 (C.I.T.). Any opinions, findings and conclusions or recommendations expressed in this material are those of the authors and do not necessarily reflect the views of the National Science Foundation. N.M.A. was supported by a National Institutes of Health (NIH) Genomics Training grant (T32 HG 000047). D.F.S. is an investigator of the HHMI and this research was supported by the HHMI.

---

**Fig. S1.**
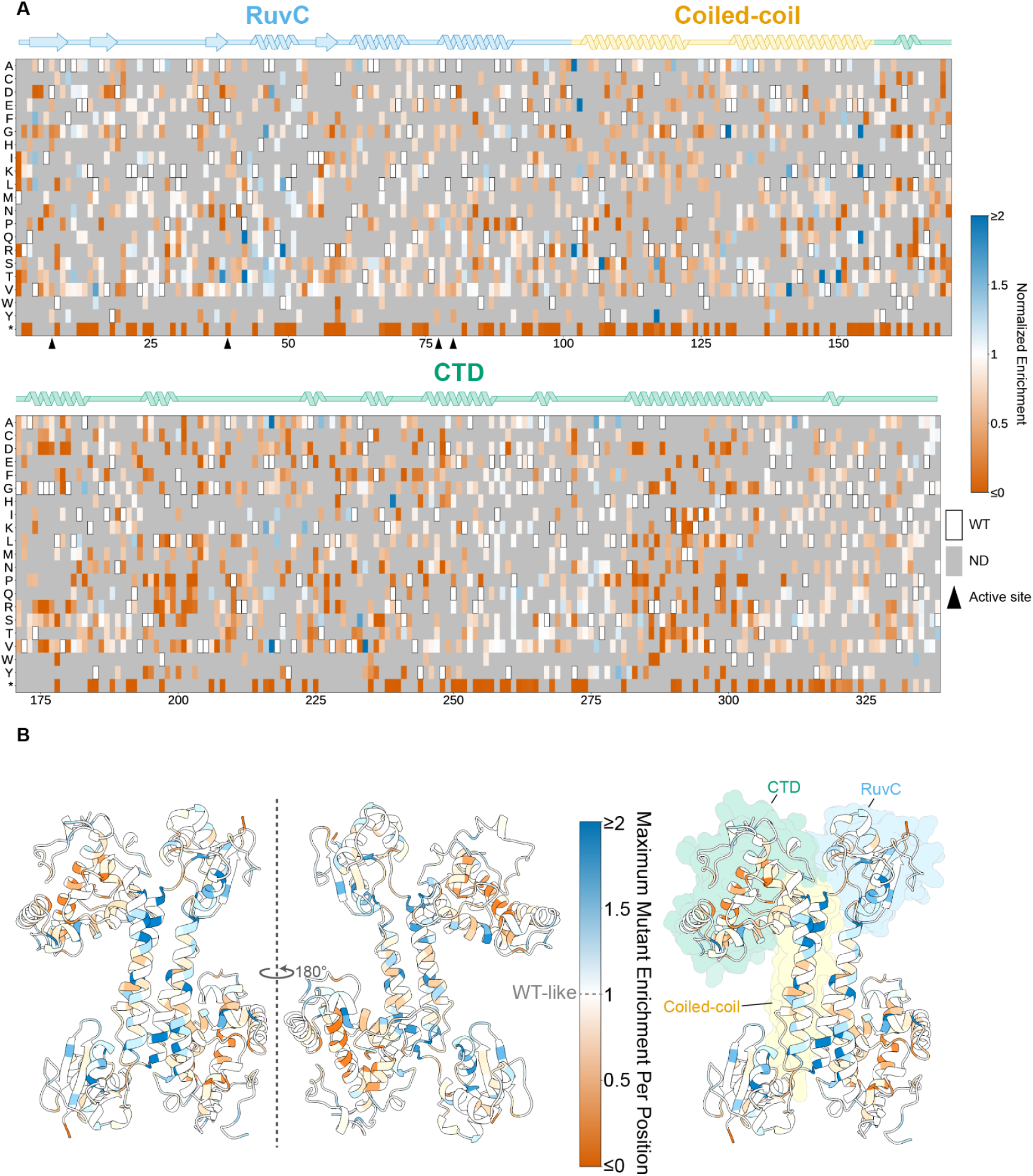
Mutation hotspots identified in epPCR library. (A) Heat map displaying enrichment values of single amino acid mutation variants within epPCR library. TasR secondary structure and domain annotations are listed above with the RuvC domain in blue, the coiled-coil dimerization domain in yellow, and the C-terminal domain (CTD) in green. Outlined white boxes denote wild-type resides. Grey boxes represent mutations with no data (ND). Black triangles indicate the active-site catalytic residues. (B) AlphaFold Apo-ParTasR model colored by maximum normalized enrichment scores at each position on one monomer and the ParTasR domain annotations imposed on the second protein monomer.

**Fig. S2.**
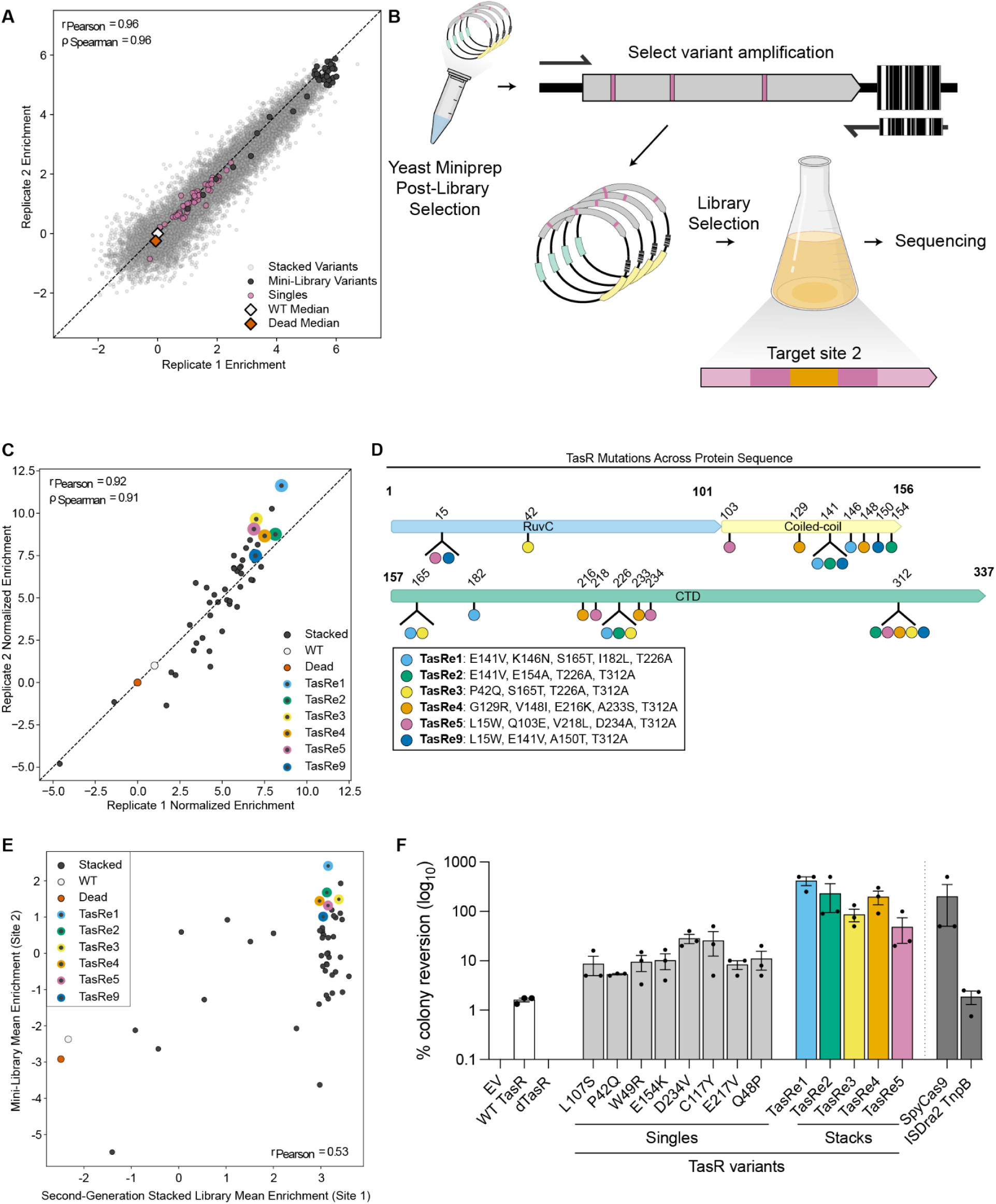
Evaluation of hit-derived variant library against secondary target. (A) The 48 variants selected from the second generation stacked library replicate correlation for assessment at an independent target site colored in charcoal. Wild-type and dead (D7A) TasR were also included in the construction of this mini-library. (B) Schematic illustrating the construction of the mini-library from the 48 selected variants recovered from the yeast minipreps of the 8 hours (h) selective (-ade) condition of the second-generation combinatorial library. Variant sequences were amplified using ssDNA oligos annealing to their corresponding known barcodes and assembled into the same yeast expression vector. The reconstructed variants were subsequently barcoded and subjected to the library selection protocol at target site 2. (C) Variant activity in the mini-library at target site 2. Enrichment scores of mini-library variants screened at target site 2 in yeast. Wild-type TasR and catalytically inactive (D7A) TasR controls are shown in white and vermillion, respectively. Six variants exhibiting high activity as both target site 1 and 2, while representing diverse mutational profiles, are highlighted with colored borders. (D) Distribution of mutations across high-activity TasR variants. Schematic illustrating the identity and position of mutations within selected high-performing combinatorial variants, mapped onto the domain architecture of TasR. Residue numbers indicate mutation positions, while colored circles denote the specific variants in which each mutation occurs. (E) Mean enrichment of stacked-mutation variants in the Site 2 mini-library plotted against their mean enrichment in the Site 1 second-generation library. Stacked-mutation variants (charcoal) represent designed combinatorial variants shared between the two libraries. WT (white circle) and dead (vermillion circle) controls are also shown. For the Site 2 mini-library, control values represent the corresponding WT and dead enrichment values within that library. For the Site 1 second-generation library, WT and dead control values represent the median enrichment across all synonymous WT barcodes and all early-stop-codon variants with ≥5 reads per replicate, respectively. (F) Activity of single and stacked mutation variants assessed by percent colony reversion with the yeast cleavage assay and compared to wild-type TasR, an empty vector (EV), and a catalytically inactive (D7A) (dead) TasR controls. Wild-type SpyCas9 and ISDra2 TnpB were included as reference editors. Data represent mean ± s.e.m. (n = 3 technical plating replicates) where variants were expressed in *S. cerevisiae* for 24 h and titer plated on selective (-ade) and non-selective (+ade) media.

**Fig. S3.**
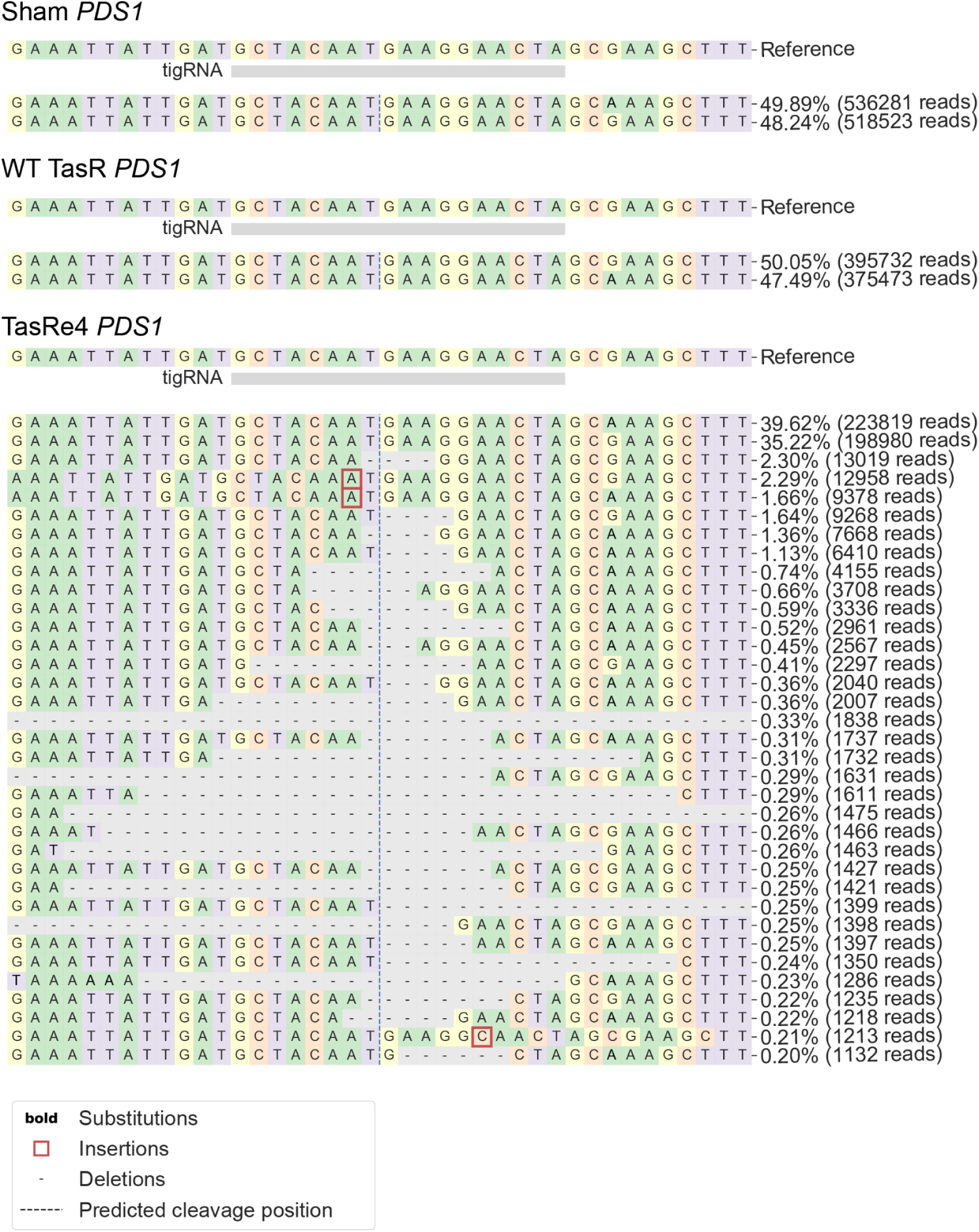
**TasR-mediated indel profiles in *N. benthamiana.*** Allele frequency tables generated by CRISPResso (v2.3.3) analysis, centered on the 18 bp tigRNA target sequence, for Sham, wild-type (WT) TasR, and TasRe4 at the single genomic locus Nb*PDS1* in *N. benthamiana.* As *N. benthamiana* is an allotetraploid, reads from both *PDS* homologs were edited and amplified hence the apparent substitutions (see Sham reads).

**Fig. S4.**
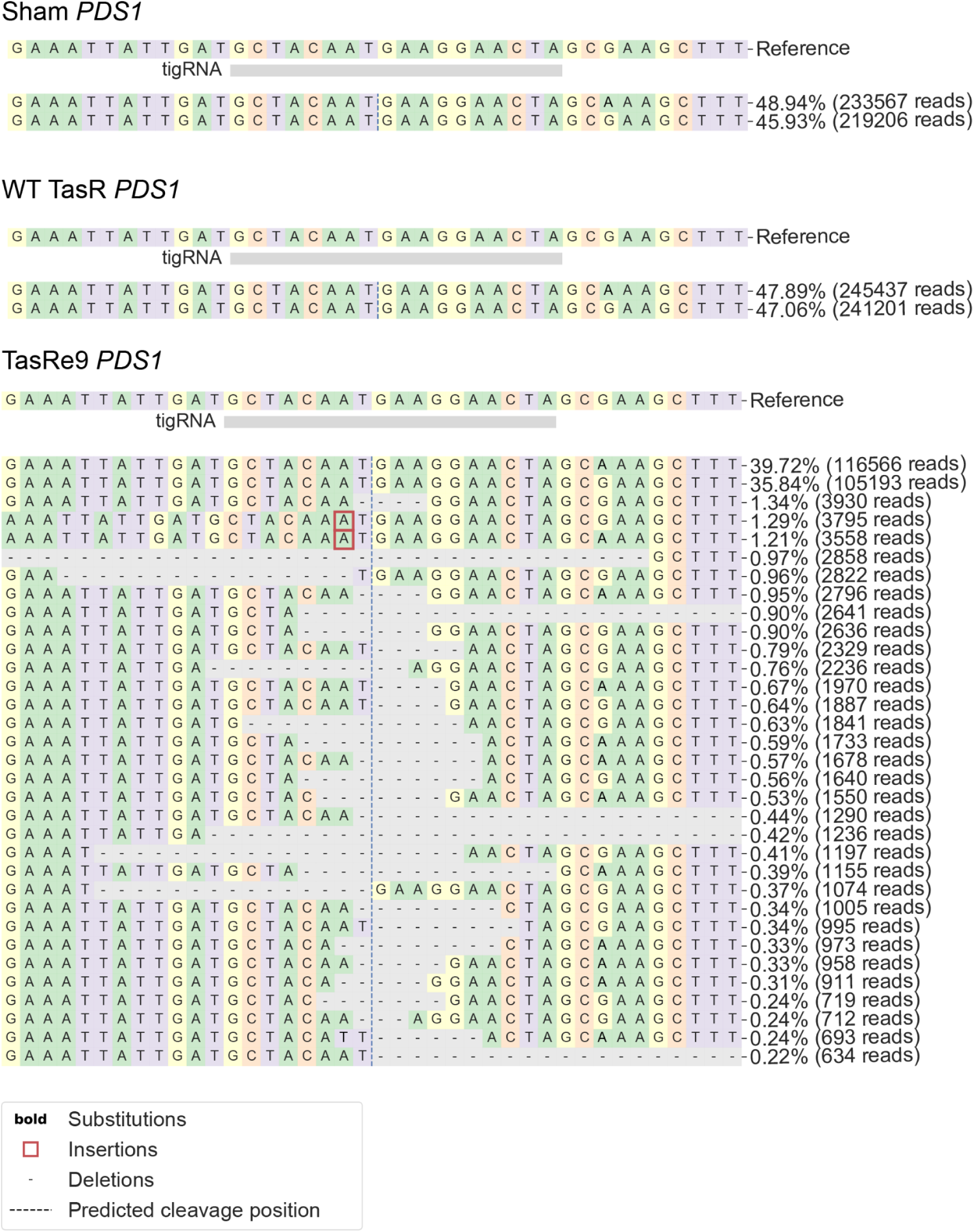
**TasR-mediated indel profiles in *N. benthamiana.*** Allele frequency tables generated by CRISPResso (v2.3.3) analysis, centered on the 18 bp tigRNA target sequence, for Sham, wild-type (WT) TasR, and TasRe9 at the single genomic locus Nb*PDS1* in *N. benthamiana.* As *N. benthamiana* is an allotetraploid, reads from both *PDS* homologs were edited and amplified hence the apparent substitutions (see Sham reads).

**Table S1.** Library Metrics

| Library type | Library synthesis method | Unique Library Variants (#) | Median # barcode per variant | Average mutation # per variant |
|---|---|---|---|---|
| **epPCR Library** | error-prone PCR | 149,230 (unfiltered); 7,697 (filtered ≥10 barcodes) | 6 (unfiltered); 10 (filtered) | 3.2 (unfiltered); 2.4 (filtered) |
| **Second-generation combinatorial library** | Nicking Mutagenesis | 13,374 | 24 | 4.15 |
| **Hit-derived mini-library** | Barcode-derived variant amplification | 50 | 17 | 4.1 |

**Table S2.**
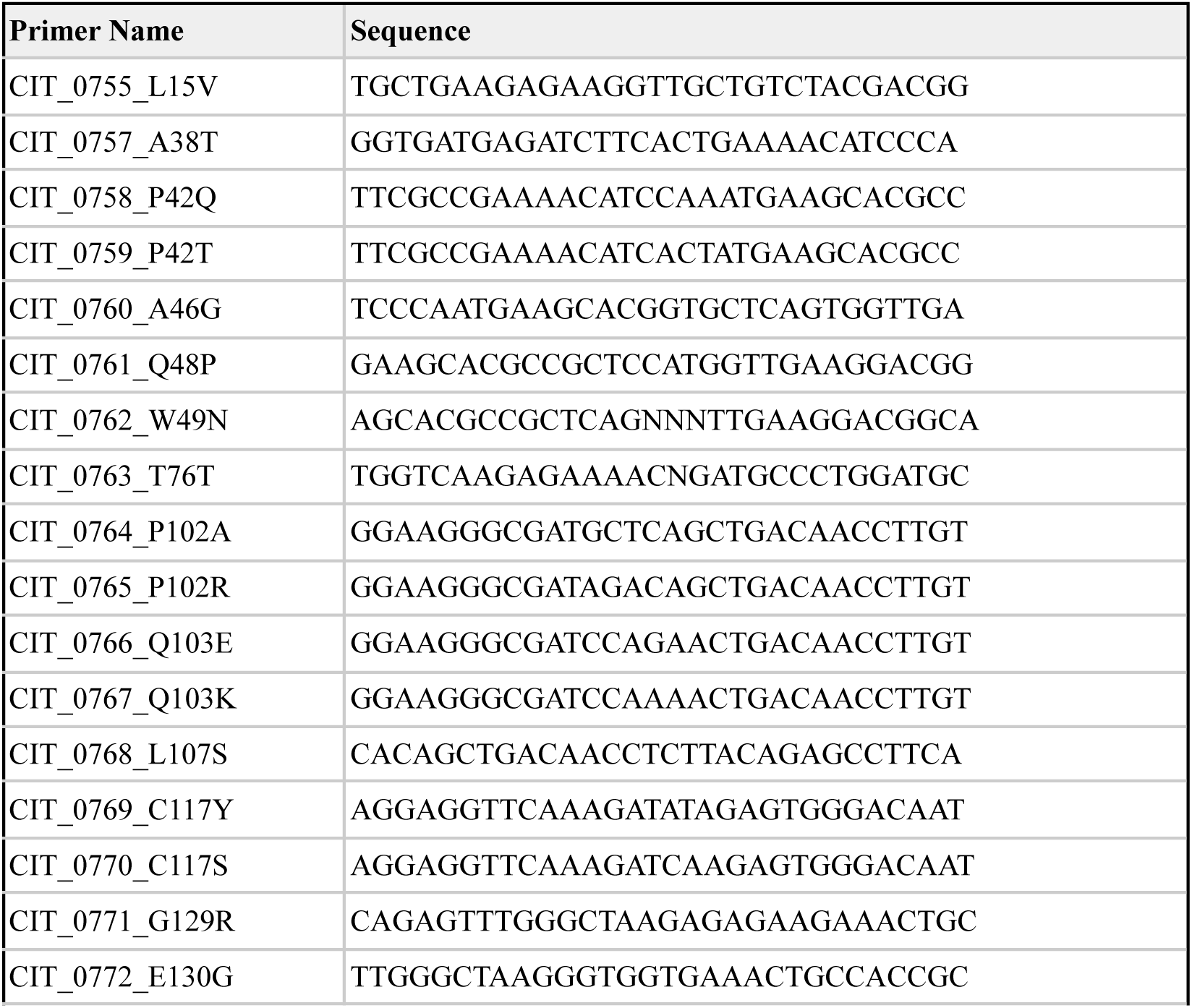
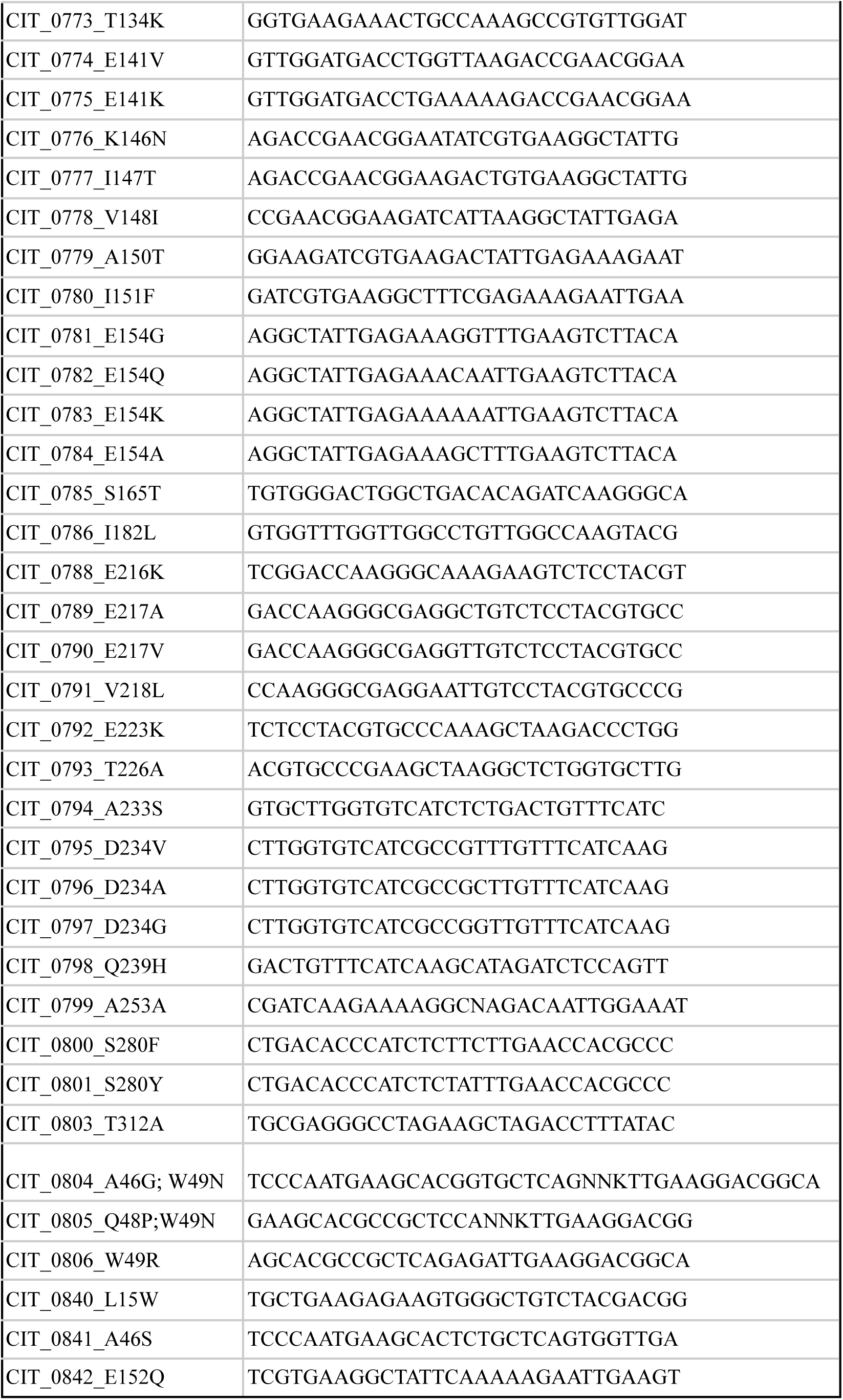
Primers used for second-generation library construction

**Table S3:**
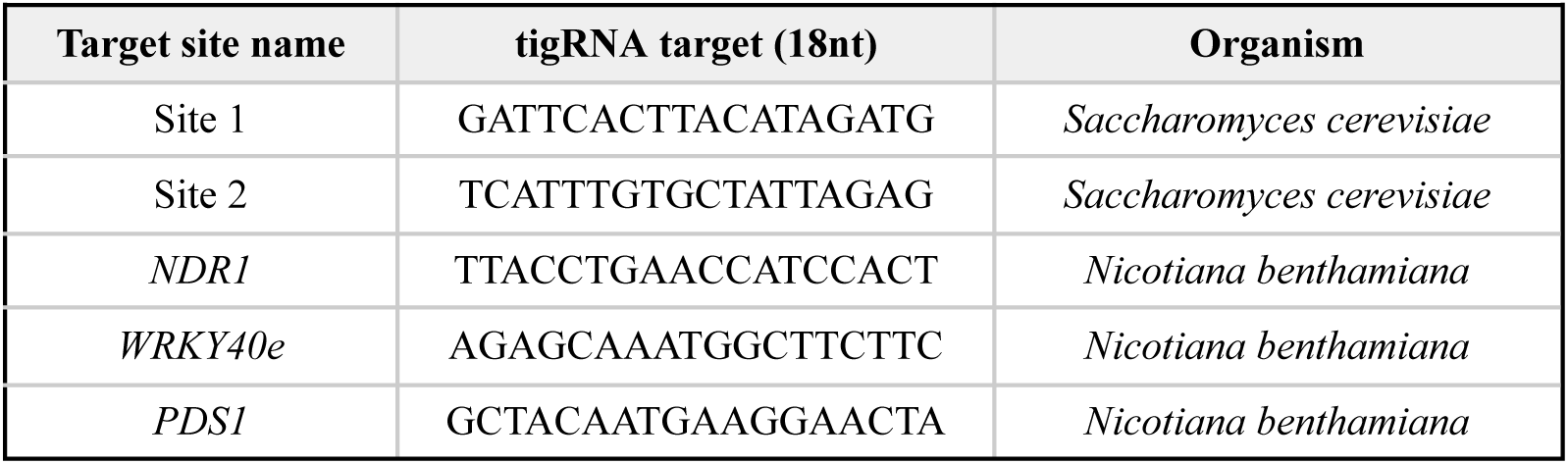
Genomic target sites for yeast screening and evaluation in plants

**Table S4:** TasR variant protein sequences

| TasR variant | Amino acid sequence |
|---|---|
| TasRe1 | MRQVAIDWAFAEEKLAVYDGKKVLKAVPKLKAGDEIFAENIPMKHAAQWLKDGIIIRRCRPNDTAALRKEIGQEKTDALDAKLIWQLAEAHPEKFREWKGDPQLTTLYRAFKEVQRCRVGQSNRVWAKGEETATAVLDDLVKTERNIVKAIEKELKSYKVWDWLTQIKGIGVATGGGLVGLLAKYGIENIRQVSSLWHLFGLHVVEGKAPRRTKGEEVSYVPEAKALVLGVIADCFIKQRSPVYRDIYDQEKARQLEIEYPEGELASKFLGYKKSDTHLSLNHAHRRAIRKMMKIFVQHVWLAWRTCEGLETRPLYCHEYLGHEHFIEPPVKIQPEGS |
| TasRe2 | MRQVAIDWAFAEEKLAVYDGKKVLKAVPKLKAGDEIFAENIPMKHAAQWLKDGIIIRRCRPNDTAALRKEIGQEKTDALDAKLIWQLAEAHPEKFREWKGDPQLTTLYRAFKEVQRCRVGQSNRVWAKGEETATAVLDDLVKTERKIVKAIEKALKSYKVWDWLSQIKGIGVATGGGLVGLIAKYGIENIRQVSSLWHLFGLHVVEGKAPRRTKGEEVSYVPEAKALVLGVIADCFIKQRSPVYRDIYDQEKARQLEIEYPEGELASKFLGYKKSDTHLSLNHAHRRAIRKMMKIFVQHVWLAWRTCEGLEARPLYCHEYLGHEHFIEPPVKIQPEGS |
| TasRe3 | MRQVAIDWAFAEEKLAVYDGKKVLKAVPKLKAGDEIFAENIQMKHAAQWLKDGIIIRRCRPNDTAALRKEIGQEKTDALDAKLIWQLAEAHPEKFREWKGDPQLTTLYRAFKEVQRCRVGQSNRVWAKGEETATAVLDDLEKTERKIVKAIEKELKSYKVWDWLTQIKGIGVATGGGLVGLIAKYGIENIRQVSSLWHLFGLHVVEGKAPRRTKGEEVSYVPEAKALVLGVIADCFIKQRSPVYRDIYDQEKARQLEIEYPEGELASKFLGYKKSDTHLSLNHAHRRAIRKMMKIFVQHVWLAWRTCEGLEARPLYCHEYLGHEHFIEPPVKIQPEGS |
| TasRe4 | MRQVAIDWAFAEEKLAVYDGKKVLKAVPKLKAGDEIFAENIPMKHAAQWLKDGIIIRRCRPNDTAALRKEIGQEKTDALDAKLIWQLAEAHPEKFREWKGDPQLTTLYRAFKEVQRCRVGQSNRVWAKREETATAVLDDLEKTERKIIKAIEKELKSYKVWDWLSQIKGIGVATGGGLVGLIAKYGIENIRQVSSLWHLFGLHVVEGKAPRRTKGKEVSYVPEAKTLVLGVISDCFIKQRSPVYRDIYDQEKARQLEIEYPEGELASKFLGYKKSDTHLSLNHAHRRAIRKMMKIFVQHVWLAWRTCEGLEARPLYCHEYLGHEHFIEPPVKIQPEGS |
| TasRe5 | MRQVAIDWAFAEEKWAVYDGKKVLKAVPKLKAGDEIFAENIPMKHAAQWLKDGIIIRRCRPNDTAALRKEIGQEKTDALDAKLIWQLAEAHPEKFREWKGDPELTTLYRAFKEVQRCRVGQSNRVWAKGEETATAVLDDLEKTERKIVKAIEKELKSYKVWDWLSQIKGIGVATGGGLVGLIAKYGIENIRQVSSLWHLFGLHVVEGKAPRRTKGEELSYVPEAKTLVLGVIAACFIKQRSPVYRDIYDQEKARQLEIEYPEGELASKFLGYKKSDTHLSLNHAHRRAIRKMMKIFVQHVWLAWRTCEGLEARPLYCHEYLGHEHFIEPPVKIQPEGS |
| TasRe9 | MRQVAIDWAFAEEKWAVYDGKKVLKAVPKLKAGDEIFAENIPMKHAAQWLKDGIIIRRCRPNDTAALRKEIGQEKTDALDAKLIWQLAEAHPEKFREWKGDPQLTTLYRAFKEVQRCRVGQSNRVWAKGEETATAVLDDLVKTERKIVKTIEKELKSYKVWDWLSQIKGIGVATGGGLVGLIAKYGIENIRQVSSLWHLFGLHVVEGKAPRRTKGEEVSYVPEAKTLVLGVIADCFIKQRSPVYRDIYDQEKARQLEIEYPEGELASKFLGYKKSDTHLSLNHAHRRAIRKMMKIFVQHVWLAWRTCEGLEARPLYCHEYLGHEHFIEPPVKIQPEGS |

